# Understanding of the 2-oxoglutarate dehydrogenase and 2-oxoadipate dehydrogenase assembly with the E2o core relevant to a hybrid complex formation

**DOI:** 10.1101/2021.02.03.429618

**Authors:** Xu Zhang, Natalia S. Nemeria, Frank Jordan

## Abstract

The 2-oxoglutarate (OG) dehydrogenase complex (OGDHc) is a key enzyme in the tricarboxylic acid cycle (TCA) and comprises multiple copies of three components: 2-oxoglutarate dehydrogenase (hE1o), dihydrolipoamide succinyltransferase (hE2o), and dihydrolipoamide dehydrogenase (hE3). The OGDHc is one of the major regulators of mitochondrial metabolism through NADH and reactive oxygen species levels and impacts cell metabolic and cell signaling pathways through the coupling of OG metabolism to gene transcription, related to tumor cell proliferation and aging. The reduced OGDHc activity is linked to a number of neurodegenerative diseases. Evidence was obtained for the formation of a hybrid 2-oxo acid dehydrogenase complex between the OGDHc and its homologue 2-oxoadipate (OA) dehydrogenase (hE1a) in the L-lysine metabolic pathway, suggesting a potential cross-talk between the two distinct metabolic pathways. These findings raised fundamental questions about assembly of hE1a and hE1o to the hE2o core. Due to the lack of an atomic structure of the OGDHc from any sources, and of knowledge about exact distribution of components around the E2 core, hydrogen/deuterium exchange (HDX-MS) and chemical cross-linking mass spectrometry (CL-MS) have been carried out in binary hE1o-hE2o, hE1a-hE2o, hE1o-hE3 and hE2o-hE3 sub-complexes followed by structural modeling. Here we report findings that revealed some similarities in the assembly of hE1o and hE1a to the hE2o core. At the same time, three regions of the hE2o core comprising residues 191-208, 273-288, and 370-386 revealed a different binding mode to hE1o and hE1a, suggesting that hE2o can differentiate between these two proteins that may have physiological consequences.

## Introduction

The 2-oxoglutarate dehydrogenase complex (OGDHc) is a key enzyme in the tricarboxylic acid cycle (TCA), comprising multiple copies of three components including 2-oxoglutarate dehydrogenase (E1o or OGDH), dihydrolipoyl succinyltransferase (E2o or DLST), and dihydrolipoyl dehydrogenase (E3 or DLD) with molecular mass of 4-10 MDa. The E1o with tightly but not covalently bound thiamin diphosphate (ThDP) decarboxylates the 2-oxoglutarate to a reactive intermediate the enamine or C2α-carbanion, which then reductively succinylates the lipoyllysyl-arm of the E2o; next the succinyl group is transferred via a transthiolesterification mechanism to CoA producing the succinyl-CoA product; finally, the E3 component reoxidizes the dihydrolipoyllysyl-E2 to lipoyllysyl-E2 producing an equivalent of NADH in the process and setting up the complex for the next turnover (1). The OGDHc represents one of the major regulators of mitochondrial metabolism through NADH and reactive oxygen species levels and impacts cell metabolic and cell signaling pathways through the coupling of the 2-oxoglutarate (OG) metabolism to the gene transcription, related to tumor cell proliferation and aging (2,3). The reduced OGDHc activity is linked to a number of neurodegenerative diseases, however, a link between reductions in the mitochondrial TCA cycle enzymes and neurodegeneration has not been established (4-7). Among the recently reported findings, is the involvement of the OGDHc in post-translational modifications (PTMs) of histones in the mammalian cell nucleus by succinylation/glutarylation that suggests a tight link between the metabolic function of OGDHc and epigenetic regulation of gene expression by PTMs (8,9). Evidence was obtained for the formation of a hybrid 2-oxo acid dehydrogenase complex between the human OGDHc (hOGDHc) and its homologue human 2-oxoadipate dehydrogenase (hE1a, also known as DHTKD1) in the L-lysine metabolic pathway, suggesting a potential cross-talk between the hOGDHc in the TCA cycle and hOADHc in L-lysine catabolism (10,11). These findings raised questions about assembly of such a hybrid complex and its physiological relevance in health and disease, especially in the light of recent genetic findings implicating the *DHTKD1* and *OGDHL* (encoding the E1o-like protein) genes in genetic etiology of several metabolic disorders (12-18) and Eosinophilic Esophagitis (EoE), a chronic allergic disorder (19). To date, there is no X-ray structure of the mammalian E1o. There are, however, X-ray structures available for the N-terminally truncated *E. coli* E1o_Δ77_ (20) and the N-terminally truncated *Mycobacterium smegmatis* α-ketoglutarate decarboxylase (*Ms*KGD_Δ115_) (21). Recently, X-ray structures were reported for the N-terminally truncated hE1a at 1.9Å (22) and at 2.25Å (23) resolution. Due to the lack of an atomic structure of the OGDHc from any sources and to the lack of knowledge about exact distribution of components around the E2 core due to their structural dynamics, hydrogen/deuterium exchange (HDX-MS) and chemical cross-linking (CL-MS) mass spectrometry have been carried out in binary hE1o-hE2o, hE1a-hE2o, hE1o-hE3 and hE2o-hE3 sub-complexes in by the Jordan group (24,25). Experiments have revealed two important facts about cross-talk between the human OGDHc (hOGDHc) in TCA cycle and human E1a (hE1a) in the L-lysine metabolic pathway: (a) The hE1o and hE1a share the same E2o and E3 components for their assembly into the corresponding OGDHc and 2-oxoadipate dehydrogenase complex (OADHc) to perform their function in two distinct metabolic pathways (10). (b) Evidence was obtained by Jordan’s group *in vitro* (10) and by Houten’s group *in vivo* (11) for the formation of a hybrid 2-oxo acid dehydrogenase complex, containing the E1o, E2o, E3 and E1a components. The notion that two E1s could share the same E2o is certainly unprecedented in the superfamily of 2-oxo acid dehydrogenase complexes and raises the fundamental questions, do hE1a and hE1o interact with the same loci of hE2o or different ones? In this paper, we addressed these questions using both experimental and computational approaches. The major conclusion from these studies is that an initial formation of the uniquely strong hE1o-hE2o and hE1a-hE2o interaction could facilitate assembly with hE3 into corresponding complexes and that hE1o and hE1a are employed the different binding loci on hE2o for the formation of the hE1a-hE2o and hE1o-hE2o sub-complexes.

## Results and Discussion

### Interactions in hE1o-hE2o and hE1a-hE2o binary sub-complexes revealed by HDX-MS

As a first step, HDX-MS experiments were conducted using the full-length hE1a and hE2o to compare and differentiate between the hE1a and hE1o binding to hE2o in binary hE1o-hE2o and hE1a-hE2o sub-complexes. Earlier, two regions from the N-terminal end of hE1o comprising residues ^18^YVEEM^22^ and ^27^LENPKSVHKSWDIF^40^ were identified as the *unique* regions which experienced a significant deuterium uptake retardation on interaction with full-length E2o, suggesting that these two predicted helical regions become less flexible on interaction with E2o (24). These results led to a major conclusion that two N-terminal regions of hE1o comprising residues ^18^YVEEM^22^ and ^27^LENPKSVHKSWDIF^40^, are likely candidates for binding with hE2o (24). On hE2o, peptide comprising residues ^144^AEAGAGKGLRSEHREKMNRMRQRIAQRLKE^173^ from the linker region and from the N-terminal region of the E2o core was suggested as critical for interaction with hE1o (24). Evidence was provided that on hE2o the binding region is extended to its core domain (24). The major conclusions from these studies are: (i) The N-terminal region of hE1o comprising residues 18-40 represents a binding locus for both hE2o and hE3. (ii) An initial formation of the hE1o-hE2o interaction facilitates assembly with hE3 to form the hOGDHc (24). In studies with hE1a by using the fluorescent labeled E2o proteins, it was demonstrated that hE1a binding to hE2o is strong enough with values of *K*_*d*_ =1.12 μM for pyrene-labeled E2o^1-173^ di-domain, comprising lipoyl domain and linker region, and *K*_*d*_ =3.98 μM for the pyrene-labeled E2o^144-386^ core domain (25). In comparison, the value of *K*_*d*_ of 0.041 μM (for hE2o^1-173^ di-domain) and *K*_*d*_ of 0.060 μM (for E2o^144-386^ core domain) were reported for hE1o (24), suggesting a different binding mode of hE1o and hE1a with the same hE2o component. Also in HDX-MS studies, it was demonstrated that the region from the N-terminal end of hE1a comprising residues 24-47 experienced significant protection from deuterium uptake in the sub-complex with hE2o, similarly to that observed for hE1o above (residues 27-40) (25). Importantly, two regions of hE1a comprising residues 485-518, which according to the recently reported X-ray structure (22) belong to the linker region connecting two halves of the hE1a monomer, and peptides in the C-terminal region of hE1a comprising residues 589-609, 646-664, and 847-874 experienced protection from deuterium uptake on interaction with hE2o (25). To probe deep inside the assembly of hE1o and hE1a to E2o, we next turned to chemical cross-linking mass spectrometry (CL-MS) to identify loci of interaction in binary E1-E2 sub-complexes.

### Loci of interaction in hE1-hE2o binary sub-complexes identified by chemical cross-linking mass-spectrometry

#### Inter-component cross links identified between hE1o and hE2o

The motivation to establish the loci of interaction in the binary sub-complexes by CL-MS was that neither the reported X-ray structures of the core domain of the E2’s (26-29), nor the single-particle cryo-EM of the hE1a-hE2o binary sub-complex (22) provided information regarding the lipoyl domain and subunit-binding region of E2o, presumed to interact with the E1o and E3 components. On the hE1o, the following cross-links were identified by using CDI (1,1’-carbonyldiimidazole, Cα-Cα distance ∼16Å to be bridged) (30) and DSBU (disuccinimidyl dibutyric urea, Cα-Cα distance ∼27Å to be bridged (31). The cross-linked Lys^31^ is located in the unique peptide comprising residues ^27^LENPKSVHKSWDIF^40^ in the N-terminal region of hE1o which experienced significant deuterium uptake retardation on interaction with hE2o (24). The other lysines are located on the hE1o surface near the ThDP and Mg^2+^-binding region (Lys^297^, Lys^309^, and Lys^362^, and Lys^495^), and in the C-terminal region (Lys^627^, Lys^657^, Lys^903^, Lys^907^ and Lys^959^) (Fig. 1, Top). The Lys^31^ in the N-terminal region of hE1o forms multiple contacts with hE2o, particularly with Lys^85^ from the linker region and Lys^286^ from the hE2o core domain in accord with HDX-MS data. On hE2o, the cross-linked Lys residues identified were located in all three domains: lipoyl domain (Lys^78^), linker region (Lys^150^) and hE2o core domain (Lys^172^, Lys^289^, and Lys^371^) (see Fig. 1 for E2o domains structure). The Lys^85^ from the linker region which forms cross-link with Lys^31^ in the N-terminal region of hE1o was identified as a ‘hot spot’, in good agreement with the OGDHc catalytic mechanism, in which the lipoyl domain serves as a swinging arm to shuttle reaction intermediates between hE1o active sites. It is not surprising that most of the cross-linked Lys residues identified on hE2o are located within the hE2o core domain, considering it carries out the principal catalytic function of hE2o, formation of succinyl-CoA or glutaryl-CoA. Particularly, Lys^371^ from the hE2o core domain, which showed interaction with the hE1o N-terminal and ThDP and Mg ^2+^-binding regions is located very near to the hE2o active center (32). In summary, the most prominent loci on hE1o interacting with hE2o is indeed the N-terminal region. On hE2o, the lysine residues from the lipoyl domain and the linker-core region form numerous cross-links with hE1o.

**Figure 1.**
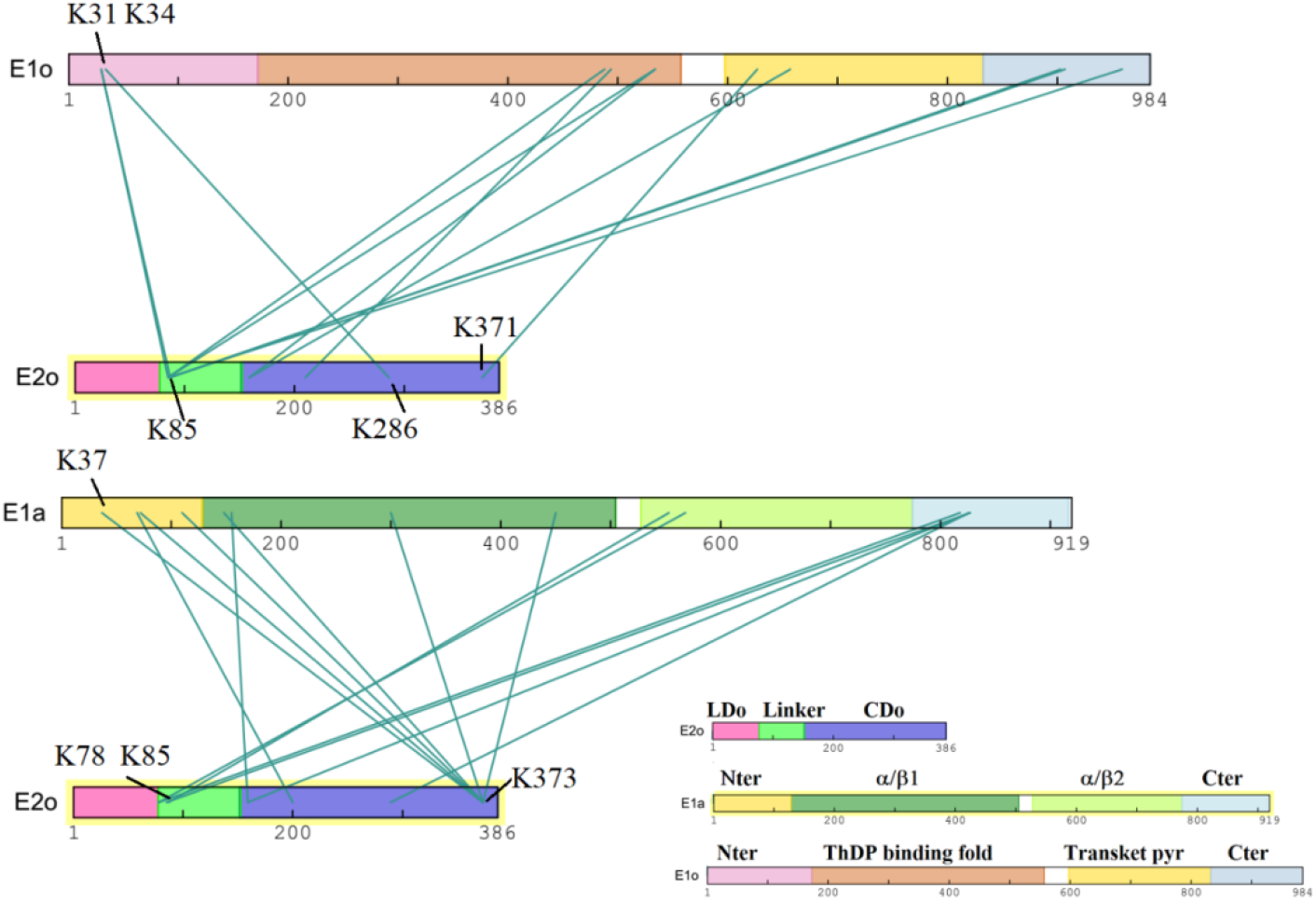
Inter-component cross-links within E1a-E2o and E1o-E2o identified with DSBU (disuccinimidyl dibutyric urea, Cα-Cα distance ∼27Å to be bridged) as cross-linker are shown in a bar plot. The bar plot was generated with xiView (45). Right-hand lower panel. Domain structure of the hE2o, hE1o, and hE1a components are marked in different colors.

#### Inter-component cross links identified between hE1a and hE2o

On the hE1a, the inter-component cross-links were identified for Lys residues from the N-terminal region (Lys^37^, Lys^148^, and Lys^188^), residues in the ThDP and Mg^2+^-binding region (Lys^300^ and Lys^450^) and those located in the C-terminal region (Lys^818^, Lys^826^/Lys^827^, and Lys^852^/Lys^854^) (25). On hE2o, the cross-linked Lys^24^, Lys^43^, Lys^66^, Lys^78^, and Lys^87^ are from the lipoyl domain; Lys^150^ and Lys^159^ are from the linker region of hE2o; and Lys^240^, Lys^286^, Lys^289^, Lys^342^, Lys^371^, and Lys^373^ are from the hE2o core domain (25). Importantly, a great number of cross-links identified were between the Lys residues from the C-terminal end of hE1a and those from the lipoyl domain of hE2o, suggesting that the C-terminal region of hE1a could be important for interaction with hE2o and for substrate channeling (25).

The interaction sites identified by CL-MS showed some similarities between these two sub-complexes. The N-terminal Lys^34^ in hE1o and Lys^37^ in hE1a both formed contacts with the hE2o core domain, but on different loci (Lys^373^ and Lys^286^, respectively). The Lys^85^ near the hE2o lipoyl domain formed cross-links with the hE1o and hE1a Lys residues from the ThDP and Mg^2+^-binding region (Lys^553^ on hE1a and Lys^534^ on hE1o) and from the C-terminal region (Lys^903^, Lys^907^, and Lys^959^ on hE1o; Lys^817^ and Lys^826^/Lys^827^ on hE1a). The major difference between the interaction loci on hE1o-hE2o and hE1a-hE2o sub-complex, is that Lys^373^ from the hE2o core domain formed multiple cross-links with Lys residues from the N-terminal region of the hE1a, however this was not observed in the hE1o-hE2o sub-complex, suggesting a different binding mode in the hE1o-hE2o and hE1a-hE2o sub-complexes.

### Structural modeling to understand the assembly in binary hE1o-hE2o and hE1a-hE2o sub-complexes

#### Three stages of structural modeling

To gain a better understanding of the hE1o-hE2o and hE1a-hE2o sub-complex assembly, the CL-MS cross-links identified were employed for structural modeling. The crystal structures of the hE1a_45-919_ at 1.9Å resolution (PDB: DHTKD1, 6sy1, ref. 22) and of the hE1a_25-919_ at 2.25 Å resolution (PDB: 6u3j ref. 23) have been reported where the N-terminal amino acids are missing. Also, there is no X-ray structure reported for the hE1o and for the full-length E2o from any sources. Under these circumstances, structural modeling was employed to understand the hE1a-hE2o and hE1o-hE2o interactions in binary sub-complexes using the protocols described in the literature (33-35). Briefly, as seen in Fig. 2, the structural modeling included three stages.

**Figure 2.**
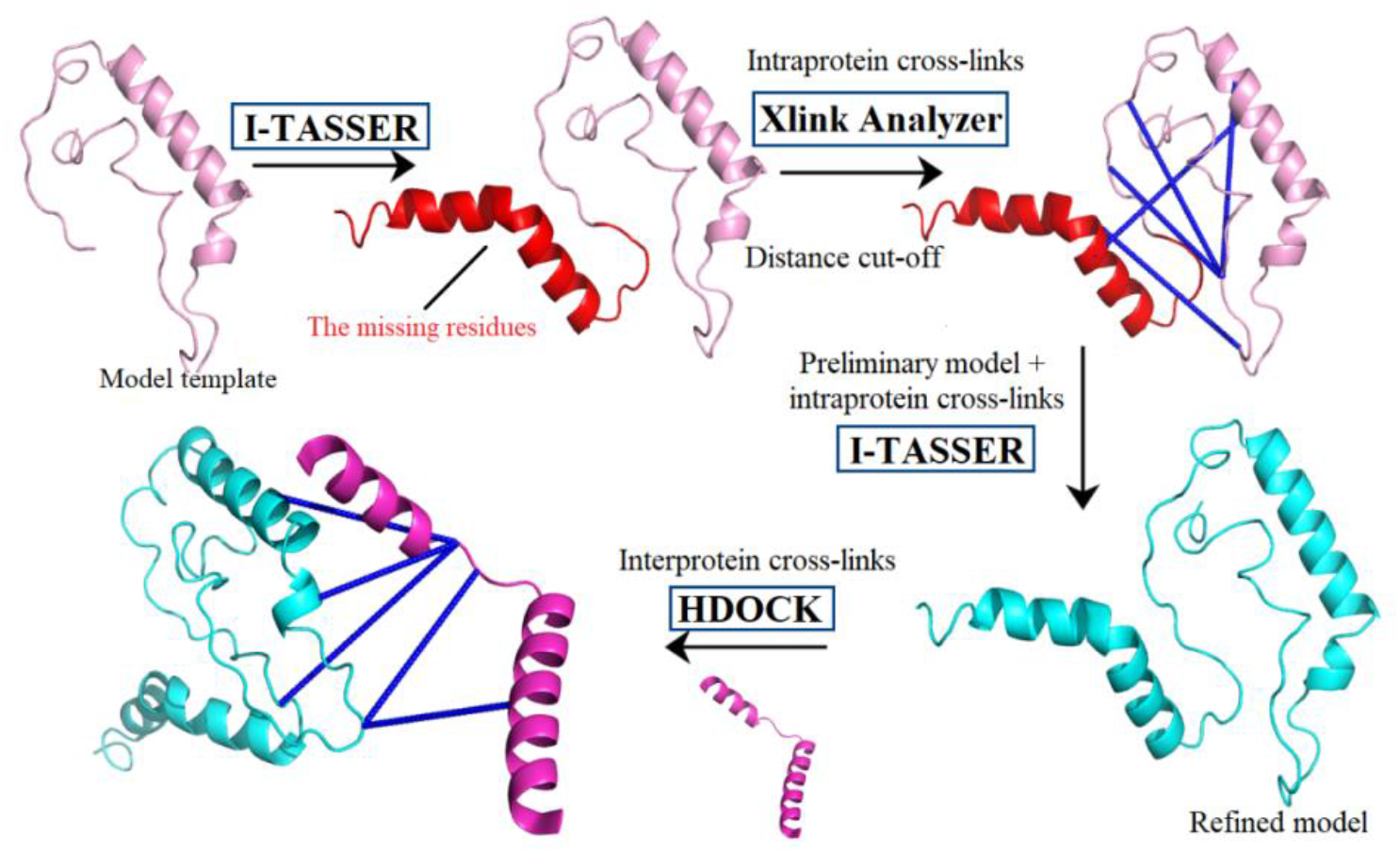
Overview of the structural modeling workflow using obtained CL-MS data. Stage 1: A preliminary model of the protein is generated using the I-TASSER algorithm (35). Stage 2: Distance cut-off is carried out to filter incompatible intra-protein (or intra-component) cross-links by using Xlink Analyzer (37). The compatible intra-protein cross-links are then used to refine the preliminary model in I-TASSER. Stage 3: The identified inter-protein (or inter-component) cross-links were used in HDOCK for protein-protein docking (38).

<u>In the first stage</u>, the preliminary model of each protein was generated by using the I-TASSER algorithm (35). For the hE1a structure, the X-ray structure of the hE1a_25-919_ (23) was used. The hE1o structure was modeled by using the reported X-ray structure of the *Ms*KGD (PDB: 2XT6; 41% identity, ref.21) as the search template. The hE2o structure was built by using multiple templates (PDB sources: 1SCZ; 6H05, ref.36; 3MAE).

<u>In the second stage</u>, the incompatible intra-protein (or intra-component) cross-links were filtered by using Xlink Analyzer (37) in visualization system UCSF Chimera (version 1.11) (www.cgi.ucsf.edu>chimera). Next, the compatible intra-protein cross-links were utilized as input with an initial model for modelling refinement, which is especially important for proteins with unknown structure, such as hE1o. The utilization of two different cross-linkers could facilitate these refinement steps. The best models of hE1o, hE1a and hE2o were selected according to the confidence score (C-score) and by how well the intra-component cross-links fit to a model. For hE2o, two models were built separately based on the hE1o-hE2o and hE1a-hE2o CL-MS data.

<u>In the third stage</u>, the resulting protein models were used as input for protein-protein docking by using the HDOCK server (38) and the docking was assisted by the inter-component cross-links identified for the hE1o-hE2o and hE1a-hE2o sub-complexes by CL-MS (24,25). Considering the length of the cross-linkers employed (2.6 Å for CDI and 12.5 Å for DSBU) and of the Lys side chains (2 × 5.5 Å), as well as the additional distance added to account for backbone dynamics (7.6-12.6 Å) (39), it was assumed that Lys residues within a Cα-Cα distance of up to 36 Å (DSBU) and 25 Å (CDI) would be preferred for the cross-linking. As was expected, some of the identified inter-component cross-links were beyond the distance cut off, hence were not considered for the next structure refining process. Cross-links identified within the Euclidean cut off mapped on the hE1o and hE2o structures are shown in Fig. 3. Also, during the docking stage, the DSBU cross-linked residues were mostly employed as distance restraints since it could provide a broader distance range for protein dynamics compared to CDI cross-linker. When the same two lysine residues were found to be cross-linked by both CDI and DSBU, the CDI’s distance restraints were employed. Protein docking was repeated with modification of different combinations of residue restraints; the HDOCK server could generate up to 100 docking solutions. The best model was selected from the top 10 solutions by taking into consideration the previous HDX-MS findings and keeping in mind our current understanding of the catalytic mechanism of the hOGDHc and hE1a: First, according to HDX-MS findings, the N-terminal regions of hE1o (residues 27-40) and of hE1a (residues 24-47) displayed significant deuterium uptake decrease upon interaction with hE2o, suggesting that the N-terminal region of both proteins could be involved in interaction with hE2o (24, 25). Second, the hE2o lipoyl domain should be near the hE1o/hE1a active center at a position such that it can transfer the succinyl/glutaryl group from hE1o/hE1a active center to the hE2o active center. In other words, if the model shows that the lipoyl domain of hE2o is too far away from the hE1o/hE1a active center, this model wouldn’t be consistent in accord with the catalytic mechanism (40). The modeled hE1o-hE2o and hE1a-hE2o sub-complex interactions are presented in Fig. 4.

**Figure 3.**
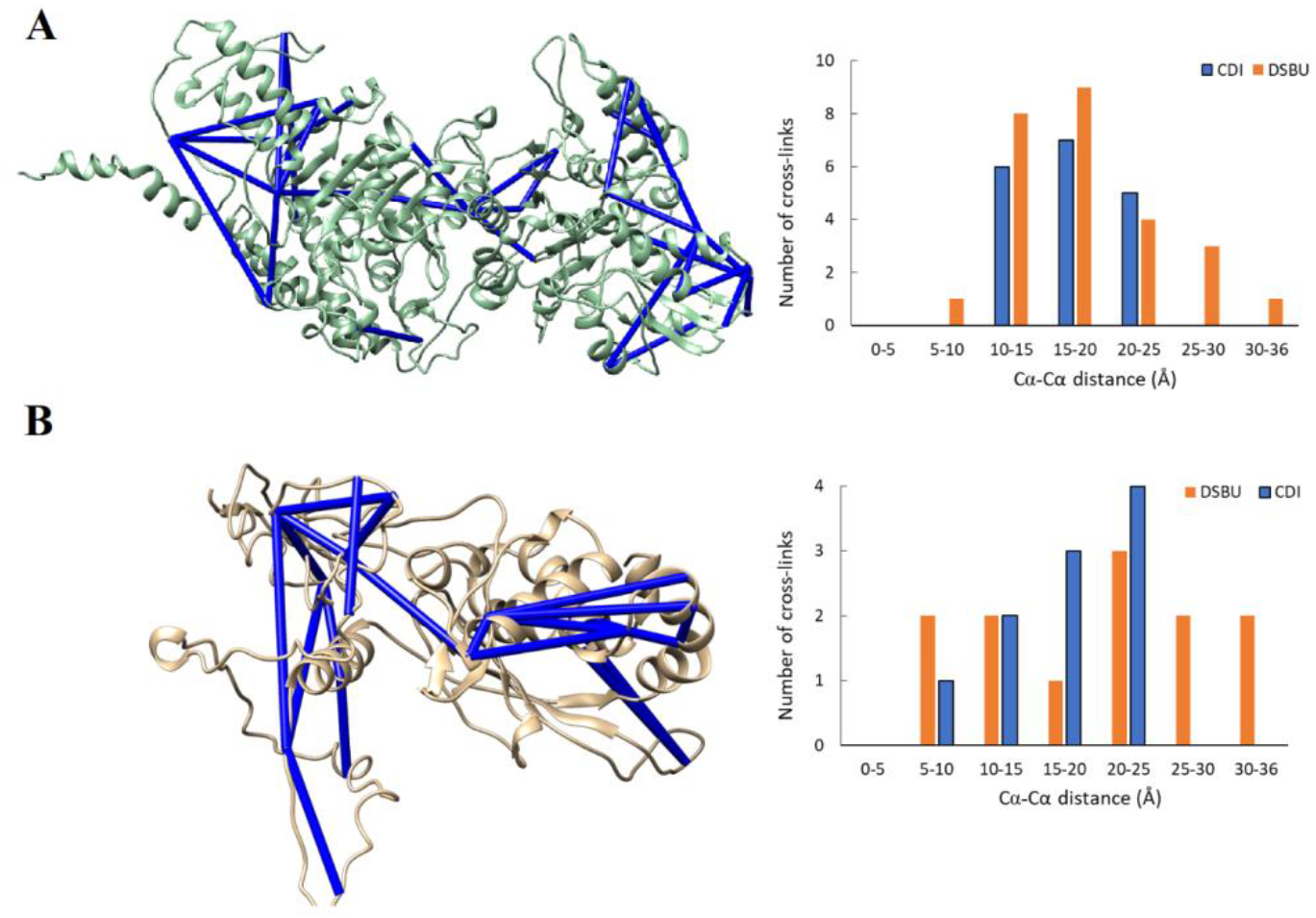
The intra-protein cross-links identified by CDI and DSBU mapped on hE1o (A) and hE2o (B) structures. (Left, top and bottom) The compatible intra-protein cross-links with false discovery rate [FDR] of <0.01 are mapped onto the modeled structures of hE1o (A) and hE2o (B) by using I-TASSER (ref). The X-ray structural templates for hE1a and hE2o are discussed in Results. (Right, top and bottom) The histogram of Cα-Cα cross-link distances for hE1o (A) and hE2o (B) revealed that all contacts are within the maximum distance span of CDI (25 Å, in *blue*) and DSBU (35 Å, in *orange*).

**Figure 4.**
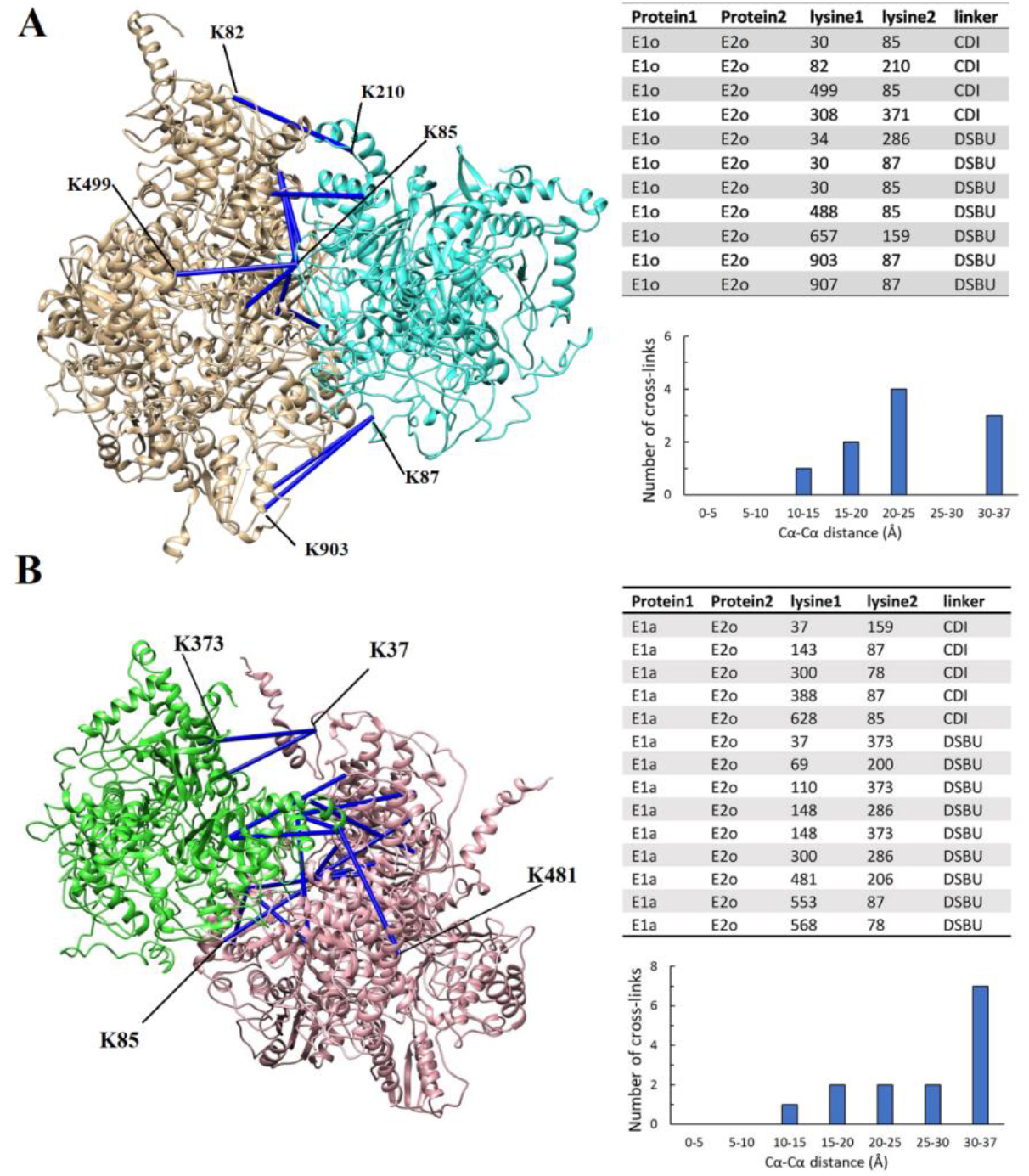
The interactions modeled for the hE1o-hE2o and hE1a-hE2o sub-complexes. In these models, the hE1o and hE1a are homodimers, and the hE2o is a heterotrimer. (Left) Interactions identified within the hE1o (in *brown*) and hE2o (in *cyan*) sub-complex (A). Interactions identified within the hE1a (in *magenda*) and hE2o (in *green*) sub-complex (B). (Right) The compatible cross-links mapped onto the hE1o-hE2o and the hE1a-hE2o sub-complexes presented in the corresponding Tables and the histograms of Cα-Cα cross-linked distances demonstrate that all contacts are within the maximum distance span of CDI (25 Å) and DSBU (36 Å).

#### Structural models for the hE1o-hE2o and hE1a-hE2o interactions

It needs to be noted that models for hE1o-hE2o and hE1a-hE2o sub-complexes share some similarities. According to models presented in Fig. 5, the hE2o binds near one of the two subunits of the hE1o/hE1a functional homodimer and ‘leans’ itself towards the active center formed at the interface between the two subunits. Particularly, the hE2o core domain is positioned on the ‘front side’ of the hE1o/hE1a with the lipoyllysyl-arm swinging around their ThDP and Mg^2+^-binding region. Structural models show that three α-helices from the hE2o core domain comprising residues 191-208, 273-289 and 370-386 are proximal to the hE1o/hE1a. Also, while the HDX-MS findings discussed above suggest that the N-terminal region of hE1o/hE1a is involved in the interactions with hE2o (becomes less open for deuterium uptake), the model studies suggest that the N-terminal end of hE1o (residues 27-40) and of hE1a (residues 24-47) are not directly involved in interaction with hE2o, but rather hidden from deuterium uptake on assembly with hE2o. At the same time, the hE2o did display different binding modes to hE1o and hE1a (Fig. 5B). In the hE1o-hE2o sub-complex, the hE2o regions from the core domain comprising residues 191-208 and 370-386 positioned closely to hE1o, while regions comprising residues 273-289 point away from hE1o. In the hE1a-hE2o sub-complex, a “clockwise twist” shifts peptide with the corresponding residues 191-280 away from the hE1a and brings the region comprising residues 273-289 closer to hE1a. At the same time, the region from the core comprising residues 370-386 from another hE2o subunit moves closer to the N-terminal end of hE1a, suggesting the different binding modes in hE1o-hE2o and hE1a-hE2o sub-complexes that allows for the hybrid complex formation. These studies begin to help us understand how hE2o could differentiate between the hE1o and hE1a and suggest further structural studies into this important enzyme complex. The cross-talk between the TCA and L-lysine degradation pathways is a novel finding with yet to be determined physiological consequences.

**Figure 5.**
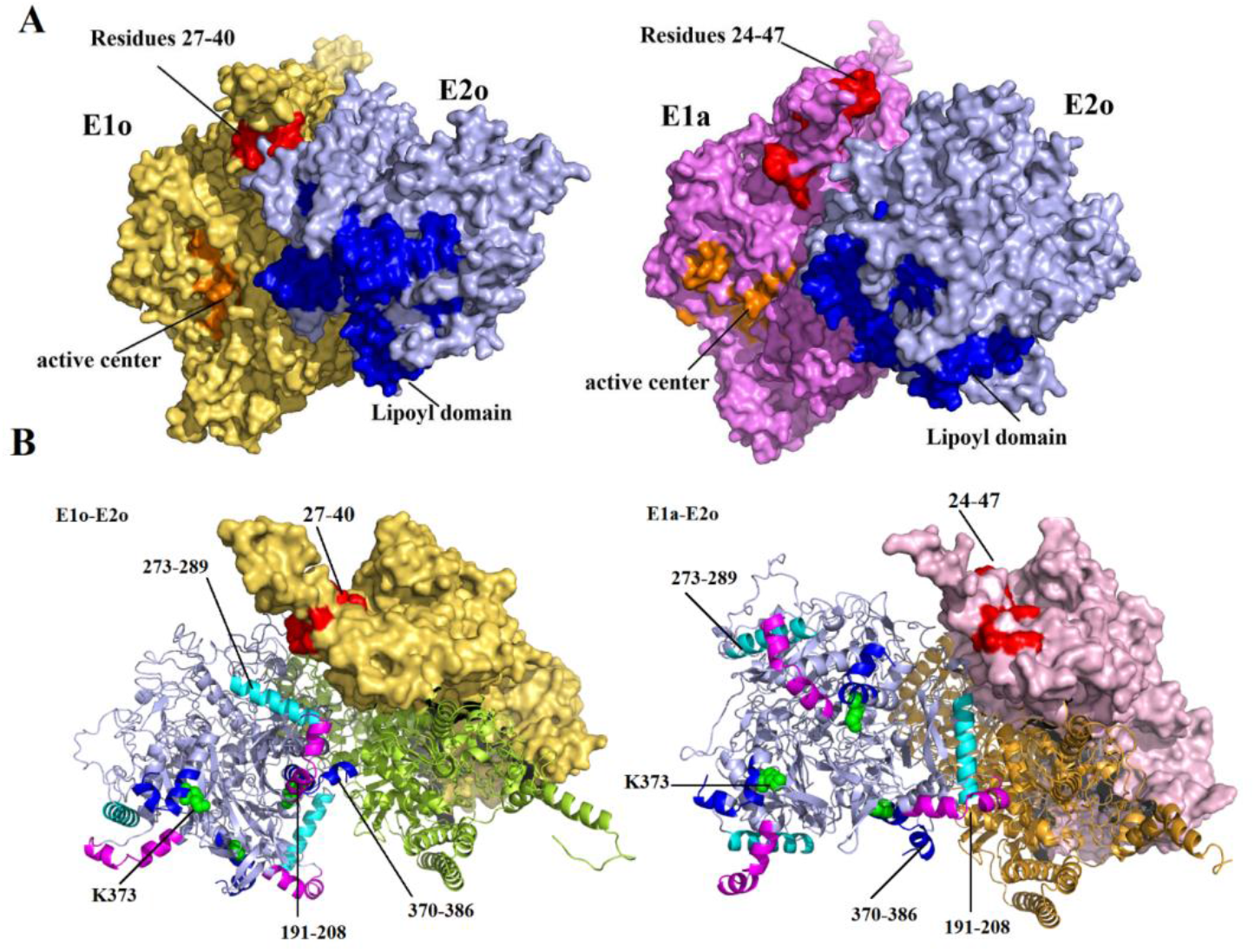
Structural models for the hE1o-hE2o and hE1a-hE2o binary sub-complexes. (A) Structural models for hE1o-hE2o (left) and hE1a-hE2o (right) sub-complexes. The proteins are colored: hE1o homodimer (in *yellow*); hE1a homodimer (in *violet*); hE2o homotrimer in both figures (in *blue*). The N-terminal residues 27-40 (hE1o) and 24-47 (hE1a) are in *red*, the E2o lipoyl domain is in *dark blue*, and the active sites of hE1o and hE1a are in *red*. (B) Different angle of viewing the models. Three regions from the E2o core domain are in different colors: residues 191-208 in *blue*, residues 273-289 in *cyan*, residues 370-386 in *purple*, Lys373 in *green*. Residues 27-40 from hE1o and residues 24-47 from hE1a are in *red*.

### Experimental Procedures

#### Chemical cross-linking mass spectrometry

Two cross-linkers, disuccinimidyldibutyric urea (DSBU) and 1,1’-carbonyldiimidazole (CDI) have been employed for cross-linking of the hE1o or hE1a with the hE2o component as follows. First, the hE1o or hE1a (1 nmol, 33 µM concentration of subunits) and hE2o (1 nmol, 33 µM concentration of subunits) were mixed in 15 µl of 20 mM K_2_HPO_4_ (pH 7.5) containing 0.5 mM ThDP, 1 mM MgCl_2_, 0.15 M NaCl and 10% glycerol. After 30 min of incubation at 20 °C, 1 µl of CDI (1 M) dissolved in DMSO was added and the cross-linking reaction was conducted at 15 °C for 45 min. When using DSBU as cross-linking reagent, 1 µl of DSBU (150 mM) dissolved in DMSO was added and the cross-linking reaction was conducted at 37 °C for 20 min. To quench the reaction, the reaction mixture was diluted to 50 µl with reaction buffer above and 1 M Tris·HCl (pH 8.0) was added to a final concentration of 20 mM. The cross-linked hE1a-hE2o were identified by SDS-PAGE (7.5%).

#### Tryptic and Glu-C proteolysis

Cross-linked samples were subjected to tryptic and Glu-C in-solution double digestion. An aliquot containing 1 nmol of the total protein was withdrawn from each reaction mixture and was placed into 70 µl of 8 M urea in 100 mM NH_4_ HCO_3_ and the samples were incubated at 60 °C. After 20 min of incubation, 2 µl of 200 mM DTT was added and the samples were incubated for an additional 40 min. Next, 3.5 µl of 200 mM iodoacetamide was added, and the samples were incubated at room temperature in the dark for 30 min. Then, 1 µl of 200 mM DTT was added, with a subsequent dilution of the reaction mixture with 0.85 ml of 100 mM NH_4_ HCO_3_. The Glu-C digestion was carried out at 50:1 protein to Glu-C (wt/wt) ratio at 37 °C. After 4 h of incubation, the reaction mixtures were subjected to tryptic digestion at 100:1 protein to trypsin (wt/wt) ratio at 37 °C. After overnight digestion, the reaction was terminated by addition of 2 µl of 95% formic acid. Digested samples were desalted on a SepPak SPE C-18 column and were dried using a SpeedVac centrifuge (Savant), then dissolved in 100 µl of 20% acetonitrile (0.05% formic acid).

#### Enrichment of cross-linked products

Peptide enrichment was carried out by using a SCX trap cartridge (Optimize tech.). The digested samples were desalted on a SepPak SPE C-18 column (Waters) and were dried in a SpeedVac (Savant), then redissolved in 100 µl 20% acetonitrile, 0.05% formic acid. The enrichment was carried out by using a SCX trap cartridge. The trap was washed with 5 volumes of 20% acetonitrile, 0.4 M NaCl, 0.05% formic acid and equilibrated with 5 volumes of 20% acetonitrile, 0.05% formic acid. Then, after loading the samples three times on the trap, the peptides were released from the trap by using a 20% acetonitrile and 0.05% formic acid with increasing concentration of NaCl: 4 volumes of 10 mM, 2 volumes of 40 mM, 4 volumes of 200 mM, and 4 volumes of 400 mM. The fractions eluted with 200 mM and 400 mM NaCl and exhibiting most of the cross-linked products were collected and concentrated in vacuo and then re-dissolved in H_2_ O containing 0.1 % formic acid.

#### Mass spectrometry

The enriched peptides were desalted by a SepPak SPE C-18 column and were dried using a SpeedVac centrifuge. Cross-linked peptides were analyzed by nano-LC-MS/MS [Dionex Ultimate 3000 RLSC nanosystem interfaced with Q Exactive HF (Thermo Fisher)]. Samples were loaded onto a self-packed 100-μm by 2-cm trap (Magic C18AQ; 5 μm and 200 Å; Michrom Bioresources, Inc.) and washed with buffer A (0.1% trifluoroacetic acid) for 5 min at a flow rate of 10 μl/min. The trap was brought in-line with the analytical column (self-packed Magic C18AQ; 3 μm and 200 Å; 75 μm by 50 cm), and peptides were eluted at 300 nl/min using a segmented linear gradient: 4 to 15% of 0.2% formic acid for 30 min; 15 to 25% of 0.16% formic acid and 80% acetonitrile for 40 min; 25 to 50% of 0.16% formic acid and 80% acetonitrile for 44 min; 50 to 90% of 0.16% formic acid; 80% of formic acid for 11 min. Data were acquired using a data-dependent acquisition procedure with a cyclic series of a full scan with a resolution of 120,000, followed by MS/MS (higher-energy C-trap dissociation; relative collision energy, 27%) of the 20 most intense ions and a dynamic exclusion duration of 20 s.

#### Analysis of cross-linking MS data

LC-MS/MS peak lists were generated using the ProteoWizard 3.0.18156 (41) software package and searched against the SwissProt database (proteome ID: UP000000558, search was done by using search GUI-3.3.117 incorporated with Peptideshaker 1.16.42 (42,43). This search showed that no contaminant protein above 5% was present in the sample. Cross-linked products were evaluated using the software tool Merox 1.6.6 (44). Only Lys-Lys cross-linked peptides were analyzed. The cross-linked peptides were manually validated, the results with False Discovery Rate (FDR) < 0.01 limit were analyzed (FDR was calculated by Merox software). Data were visualized using the xiView website (45).

#### Structural modeling

Structural modeling of the hE1a-hE2o and hE1o-hE2o interactions in binary sub-complexes was performed using the protocols described in the literature (46, 47). Briefly, in the first stage of the modeling, I-TASSER (35) was used to construct structures of the individual hE1a, hE1o and hE2o proteins. The structure of the hE1a monomer was built by using the X-ray structure of the hE1a reported recently (22, 23). The hE1o structure was modeled using the reported *Ms*KGD structure (PDB: 2XT6; 41% identity, Ref 48) as the search template. The hE2o structure was built by using multiple templates with I-TASSER (PDB: 1SCZ; 6H05; 3MAE). In the second stage, the incompatible intra-component cross-links were filtered by using the XLink Analyzer (37) in visualization system UCSF Chimera (version 1.11) (49) by applying the distance threshold of 36 Ǻ for DSBU and of 25 Ǻ for CDI. The Euclidean cut off was calculated as the sum of the length of the two extended lysine chains (2 × 5.5 Å) plus the spacer length (2.6 Å for CDI and 12.5 Å for DSBU), with an additional 7.6-12.6 Å allowed for conformational dynamics (39). The peptides identified by DSBU and CDI cross-links were mapped into the corresponding protein structures. Compatible cross-links were used for protein structure refinement in I-TASSER as assigned distance restraints. The structures of the hE1a homodimer and the hE2o homotrimer were created through alignment in PyMOL (v2.2.1) to other subunits as given in the templates. The resulting structures were used as input for protein-protein docking using the HDOCK server (38) and the docking was assisted by the inter-component cross-links obtained for the hE1a-hE2o and hE1o-hE2o sub-complex by CL-MS.

## Data availability

The mass spectrometry proteomics data have been deposited to the ProteomeXchange Consortium (www.proteomexchange.org) via the PRIDE partner repository with the dataset identifier PXD017792 and PXD023525.

## Acknowledgements

The authors acknowledge Roman Brukh at Rutgers University Newark, NJ for the help with data collection using FT-MS instrument and researchers at Rutgers University New Brunswick, NJ for the help with analyses of the cross-linked peptides by using nano-LC-MS/MS.

## Author Contributions

All three coauthors conceived the project. XZ carried out the preparation for and execution of the mass spectrometry experiment. NSN created all of the proteins. All three authors collaborated on writing the manuscript.

## Abbreviations

OGDHc: 2-oxoglutarate dehydrogenase complex
h: human
OADHc: 2-oxoadipate dehydrogenase complex
E1o: 2-oxoglutarate dehydrogenase
E1a: 2-oxoadipate dehydrogenase
E2o: dihydrolipoamide succinyltransferase
E3: dihydrolipoamide dehydrogenase
LDo: lipoyl domain
CDo: E2o catalytic domain
PTMs: post-translational modifications
TCA cycle: tricarboxylic acid cycle
ThDP: thiamin diphosphate
*DHTKD1*: gene encoding 2-oxoadipate dehydrogenase
HDX-MS: hydrogen-deuterium exchange mass spectrometry
CL-MS: chemical cross-linking mass spectrometry
FT-MS: Fourier transform mass spectrometry
cryo-EM: cryogenic electron microscopy
CDI: 1,1’-carbonyldiimidazole cross-linker
DSBU or BuUrBu: disuccinimidyl dibutyric urea
PDB: Protein Data Bank.

## References

1. Jordan, F. (2003) Current mechanistic understanding of thiamin diphosphate-dependent enzymatic reactions. Nat. Prod. Rep. 20, 184–201

2. Vatrinet, R., Leone, G., De Luise, M., Girolimetti, G., Vidone, M., Gasparre, G., Porcelli, A. M. (2017) The α-ketoglutarate dehydrogenase complex in cancer metabolic plasticity. Cancer & Metabolism 5:3. DOI 10.1186/s40170-017-0165-0

3. Burr, S. P., Costa, A. S. H., Grice, G. L., Timms, R. T., Lobb, I. T., Freisinger, P., Dodd, R. B. Dougan, G., Lehner, P. J., Frezza, C., Nathan, J. A. (2016) Mitochondrial protein lipoylation and the 2-oxoglutarate dehydrogenase complex controls HIF1α stability in aerobic conditions. Cell Metab. 25, 740–752

4. Gibson, G. E., Park, L. C. H., Sheu, K.-F.R., Blass J. P., Calingasan, N. Y. (2000) The α-ketoglutarate dehydrogenase complex in neurodegeneration. Neurochem. Intern., 36, 97–112

5. Gibson, G. E., Kingsbury, A. E., Xu, H., Lindsay, J. G., Daniel, S., Foster, O. J., Lees, A. J., Blass, J. P. (2003) Deficits in a tricarboxylic acid cycle enzyme in brains from patients with Parkinson’s disease. Neurochem. Int. 43, 129–135

6. Gibson, G. E., Xu, H., Chen, H.-L., Chen, W., Denton, T., and Zhang, S. (2015) Alphaketoglutarate dehydrogenase complex-dependent succinylation of proteins in neurons and neuronal cell lines. J. Neurochem. 134, 86–96

7. Chen, H., Denton, T. T., Xu, H., Calingasan, N., Beal, M. F., Gibson, G. E. (2016) Reductions in the mitochondrial enzyme α-ketoglutarate dehydrogenase complex in neurodegenerative disease beneficial or detrimental? J. Neurochem., 139, 823–838

8. Bao, X., Liu, Z., Zhang, W., Gladysz, K., Fung, Y. M. E., Tian, G., Xiong, Y., Wong, J. W. H., Yuen, K. W. Y., Li, X. D. (2019) Glutarylation of histone H4 Lysine 91 regulates chromatin dynamics. Mol. Cell 76, 660–675

9. Wang, Y., Guo, Y. R., Liu, K., Yin, Z., Liu, R., Xia, Y., Tan, L., Yang, P., Lee, J. H., Li, X., Hawke, D., Zheng, Y., Qian, X., Lyu, J., He, J., Xing, D., Tao, Y. J., Lu, Z. (2017) KAT2A coupled with the α-KGDH complex acts as a histone H3 succinyltransferase. Nature, 552, 273–277

10. Nemeria, N. S., Gerfen, G., Reddy Nareddy, P., Yang, L., Zhang, X., Szostak, M., Jordan, F. (2018) The mitochondrial 2-oxoadipate and 2-oxoglutarate dehydrogenase complexes share their E2 and E3 components for their function and both generate reactive oxygen species. Free Radic. Biol. Med. 115, 136–145

11. Leandro, J., Dodatko, T., Aten, J., Nemeria, N.S., Zhang, X., Jordan, F., Hendrickson, R. C., Sanchez, R., Yu, C., DeVita, R. J., Houten, S.M. (2020) DHTKD1 and OGDH display substrate overlap in cultured cells and form a hybrid 2-oxo acid dehydrogenase complex in vivo. Hum. Mol. Genet. 29, 1168–1179

12. Danhauser, K., Sauer, S., Haack, T. B., Wieland T., Staufner, C., Graf, E., Zschocke, J., Strom, T. M., Traub, T., Okun, J. G., Meitinger, T., Hoffmann, G. F., Prokisch, H., Kölker, S. (2012) DHTKD1 mutations cause 2-aminoadipic and 2-oxoadipic aciduria. Am. J. Human Gen. 91, 1082–1087

13. Stiles, A. R., Venturoni, L., Mucci, G., Elbalalesy N, Woontner, M., Goodmann, S. and Abdenur J. E. (2015) New cases of DHTKD1 mutations in patients with 2-ketoadipic aciduria. JIMD Reports 25, 15–19

14. Hagen, J., Brinke, H., Wanders, R. J. A., Knegt. A. C., Oussoren, E., Hoogeboom, A. J. M., Ruijter, G. J. G., Becker, D., Schwab, K. O., Franke, I., Duran, M., Waterham, H. R., Sass, J. O., Houten, S. M. (2015) Genetic basis of alpha-aminoadipic and alpha-ketoadipic aciduria. J. Inherit.Metab. Dis. 38, 873–879

15. Xu, W.Y., Gu, M. M., Sun, L. H., Guo, W.T., Zhu, H. B., Ma, J. F., Yuan, W.T. Kuang, Y., Ji, B. J., Wu, X. L., Chen, Y., Zhang, H. X., Sun, F. T., Lei, W. H., Chen, H. S., Wang, Z. G. (2012) A nonsence mutation in DHTKD1 causes Charcot-Marie-Tooth type 2 in a large chinese pedigree. Am. J. Hum. Genet. 91,1088–1094

16. Xu, W., Zhu, H., Gu, M., Luo, Q., Ding, J., Yao,Y., Chen, F., Wang, Z. (2013) DHTKD1 is essential for mitochondrial biogenesis and function maintenance. FEBS Letters 587, 3587–3592

17. Xu W.Y.., Zhu, H., Shen, Y., Wan, Y. H., Tu, X. D., Wu, W. T., Tang, L., Zhang, H. X., Lu, S. Y., Jin, X. L., Fei, X. L., Wang, Z. G. (2018) DHTKD1 deficiency causes Charcot-Marie-Tooth disease in mice. Mol. Cell. Biol. 38 https://doi.org/10.1128/MCB.00085-18

18. Biagosch, C., Ediga, R. D., Hensler, S. V., Faerberboeck, M., Kuehn, R., Wurst, W., Meitinger, T., Kölker, S., Sauer, S., Prokisch, H. (2017) Elevated glutaric acid levels in Dhtkd1-/Gcdh-double knockout mice challenge our current understanding of lysine metabolism. BBA Mol. Basis of disease,1863, 2220–2228

19. Sherrill, J. D., Kc, K., Wang, X., Wen, T., Chamberlin, A., Stucke, E. M., Collins, M. H., Abonia, J.P., Peng, Y., Wu, Q., Putnam, P. E., Dexheimer, P. J., Aronow, B. J., Kottyan, L. C., Kaufman, K. M., Harley, J. B., Huang, T., and Rothenberg, M. E. (2018) Whole-exome sequencing uncovers oxidoreductases DHTKD1 and OGDHL as linkers between mitochondrial disfunction and eosinophilic esophagitis. JCI Insight, 3(8) https://doi.org/10.1172/jci.insight.99922

20. Frank, R. A., Price, A. J., Northrop. F. D., Perham, R. N., Luisi, B. F. (2007) Crystal structure of the E1 component of the Escherichia coli 2-oxoglutarate dehydrogenase multienzyme complex. J. Mol. Biol. 368, 639–651

21. Wagner, T., Bellinzoni, M., Wehenkel, A., O’Hare, H. M., Alzari, P. M. (2011) Functional plasticity and allosteric regulation of α-ketoglutarate decarboxylase in central mycobacterium metabolism. Chemistry & Biology, 18, 1011–1020

22. Bezerra, G. A., Foster, W. R., Bailey, H. J., Hicks, K. G., Sauer, S. W., Dimitrov, B., McCorvie, T. J., Okun, J. G., Rutter, J., Kölker, S., Yue, W. W. (2020) Crystal structure and interaction studies of human DHTKD1 provide insight into a mitochondrial megacomplex in lysine catabolism. IUCrJ, 7, 693–706

23. Leandro J., Khamrui, S., Wang, H., Suebsuwong, C., Nemeria, N., Huynh, K., Moustakim, M., Secor, C., Wang, M., Dodatko, T., Stauffer, B., Wilson, C. G., Yu, C., Arkin, M. R., Jordan, F., Sanchez, R., DeVita, R. J., Lazarus, M. B., Houten, S. (2020) Inhibition and crystal structure of the human DHTKD1-Thiamin diphosphate complex. ACS Chem. Biol. 15, 2041–2047

24. Zhou, J., Yang, L., Ozohanics, O., Zhang, X., Wang, J., Ambrus, A., Arjunan, P., Brukh, R., Nemeria, N. S., Furey, W., Jordan, F. (2018) A multipronged approach unravels unprecedented protein-protein interactions in the human 2-oxoglutarate dehydrogenase multienzyme complex. J. Biol. Chem., 293, 13204–13213

25. Zhang, X., Nemeria, N. S., Leandro, J., Houten, S., Lazarus, M. B., Gerfen, G. J., Ozohanics, O., Ambrus, A., Nagy, B., Brukh, R., Jordan, F., (2020) Structure-function analyses of the G729R 2-oxoadipate dehydrogenase genetic variant associated with L-lysine metabolism disorder. J. Biol. Chem. 295, 8078–8095

26. DeRosier, D. J., Oliver, R. M. & Reed, L. J. (1971) Crystallization and preliminary structural analysis of dihydrolipoyltranssuccinylase, the core of the 2-oxoglutarate dehydrogenase complex. Proc. Natl. Acad. Sci. USA, 68, 1135–1137

27. Knapp, J. E., Mitchell, D. T., Yazdi, M. A., Ernst, S. R., Reed, L. J., and Hackert, M. L. (1998) Crystal structure of the truncated cubic core component of the Escherichia coli 2 oxoglutarate dehydrogenase multienzyme complex. J. Mol. Biol. 280, 655–668

28. Knapp, J. E., Carroll, D., Lawson, J. E., Ernst, S. R., Reed, L. J., and Hackert, M. L. (2000) Expression, purification, and structural analysis of the trimeric form of the catalytic domain of the Escherichia coli dihydrolipoamide succinyltrancferase. Protein Sci. 9, 37–48

29. Suzuki, K., Adachi, W., Yamada, N., Tsunoda, M., Koike, K., Koike, M., and Sekénaka, A. (2002) Crystallization and preliminary X-ray analysis of the full-size cubic core of pig 2-oxoglutarate dehydrogenase complex. Acta Crystallographica Section D, Structural Biology, 58, 833–835

30. Hage, C., Iacobucci, C., Rehkamp, A., Arlt, C., Sinz, A. (2017) The first zero-length mass spectrometry -cleavable cross-linker for protein structure analysis. Angew. Chem. Int. Ed. 56, 14551 – 14555

31. Müller, M.Q., Dreiocker, F., Ihling, C.H., Sch໤fer, M., Sinz, A. (2010) Cleavable cross-linker for protein structure analysis: reliable identification of crosslinking products by tandem MS. Anal. Chem. 82, 6958–6968

32. Chakraborty, J., Nemeria, N. S., Farinas, E. T., Jordan F. (2018) Catalysis of transthioacylation in the active centers of dihydrolipoamideacyltransacetylase components of 2-oxo acid dehydrogenase complexes. FEBS Open Bio. 8, 880–896

33. Iacobucci, C., Götze, Michael., Ihling, C. H., Piotrowski, C., Arlt, C., Schäfer, M., Hage, C., Schmidt, R., Sinz, A. (2018) A cross-linking/mass spectrometry workflow based on MS-cleavable cross-linkers and the MeroX software for studying protein structures and protein-protein interactions. Nature Protocols 13, 2864–2889

34. Klykov, O., Steigenberger, B., Pektas, S., Fasci, D., Heck, A. J. R., Scheltema, R. A. (2018) Efficient and robust proteome-wide approaches for cross-linking mass spectrometry. Nature Protocols 13, 2964–2990

35. Roy, A., Kucukural, A., Zhang, Y. (2010) I-TASSER: a unified platform for automated protein structure and function prediction. Nature Protocols 5, 725–738

36. Tanina, A., Wohlkonig, A., Soror, S. H., Flipo, M., Villemagne, B., Prevet, H., Deprez, B., Moune M., Peree, H., Meyer, F., Baulard, A. R., Willand, N., Wintjens, R. (2018) A comprehensive analysis of the protein-ligand interactions in crystal structures of Mycobacterium tuberculosis EthR. Biochim. Biophys. Acta Proteins Proteom 1867, 248–258

37. Kosinski, J., von Appen, A., Ori, A., Karius, K., Muller, C.W., Beck, M. (2015). Xlink analyzer software for analysis and visualization of cross-linking data in the context of three-dimensional structures. J. Struct. Biol. 189,177–183

38. Yan, Y., Tao, H., He, J., Huang. S.-Y. (2020) The HDOCK server for integrated protein-protei docking. Nature Protocols 15, 1829–1852

39. Kahraman, A., Herzog, F., Leitner, A., Rosenberger, G., Aebersold, R., Malmström, L. (2013) Cross-link guided molecular modeling with ROSETTA. PloS One. 8, e73411

40. Pan, K., Jordan, F. (1998) D, L-S-methyllipoic acid methyl ester, a kinetically viable model for S-protonated lipoic acid as the oxidizing agent in reductive acyl transfers catalyzed by the 2-oxoacid dehydrogenase multienzyme complexes. Biochemistry 37, 1357–1364

41. Chambers, M.C., MacLean, B., Burke, R., Amode, D., Ruderman, D.L., Neumann, S., Gatto, L., Fischer, B., Pratt, B., Egertson, J., Hoff, K., Kessner, D., Tasman, N., Shulman, N., Frewen, B., Baker, T.A., Brusniak, M.-Y., Paulse, C., Creasy, D., Flashner, L., Kani, K., Moulding, C., Seymour, S.L., Nuwaysir, L.M., Lefebvre, B., Kuhlmann, F., Roark, J., Rainer, P., Detlev, S., Hemenway, T. Huhmer, A., Langridge, J., Connolly, B., Chadick, T., Holly, K., Eckels, J., Deutsch, E.W., Moritz, R.L., Katz, J.E., Agus, D.B., MacCoss, M., Tabb, D. L., and Mallick, P. A. (2012) A cross-platform toolkit for mass spectrometry and proteomics. Nature Biotechnol. 30, 918–920

42. Barsnes, H., Vaudel, M. (2018) SearchGUI: a highly adaptable common interface for proteomics search and de novo engines. J. Proteome Res. 17, 2552–2555

43. Vaudel, M., Burkhart, J. M., Zahedi, R. P., Oveland, E., Berven, F. S., Sickmann, A., Martens, L., Barsnes, H. (2015) PeptideShaker enables reanalysis of MS-derived proteomics data sets. Nature Biotechnol. 33, 22–24

44. Gotze, M., Pettelkau, J., Fritzsche, R., Ihling, C. H., Schäfer, M., and Sinz, A. (2015) Automated assignment of MS/MS cleavable cross-links in protein 3D-structure analysis. J. Am. Soc. Mass 1. Spectrom. 26, 83–97

45. Combe, C. W., Fischer, L., Rappsilber, J. (2015) xiNET: cross-link network maps with residue resolution. Molecular & Cellular Proteomics 14, 1137–1147

46. Orbán-Németh, Z., Beveridge, R., Hollenstein, D. M., Rampler, E., Stranzl, T., Hudecz, O., Doblmann, J., Schlögelhofer, P., Mechtler, K. (2018) Structural prediction of protein models using distance restraints derived from cross-linking mass spectrometry data. Nature Protocols 13, 478–494

47. Fux, A., Korotkov, V. S., Schneider, M, Antes, I., Sieber S. A. (2019) Chemical cross-linking enables drafting ClpXP proximity maps and taking snapshots of in situ interaction networks. Cell Chem. Biol. 26, 48-59.e7

48. Wagner, T., Barilone, N., Alzari, P. M., Bellinzoni, M. (2014) A dual conformation of the pos decarboxylation intermediate is associated with distinct enzyme states in mycobacterial KGD (α-ketoglutarate decarboxylase). Biochem. J., 457, 425–434

49. Pettersen, E. F., Goddard, T. D., Huang, C. C., Couch, G. S., Greenblatt, D. M., Meng, E. C., Ferrin, T.E. (2004) UCSF chimera - a visualization system for exploratory research and analysis. J. Comput. Chem. 25, 1605–1612

